# Genetic Code Expansion, Enzymatic Modification, and C-Terminal Labeling Enable Facile Production of Highly Modified α-Synuclein

**DOI:** 10.64898/2026.06.24.734353

**Authors:** Bernard Abakah, Marie Shimogawa, Paola Miranda-Castrodad, Elizabeth Rhoades, E. James Petersson

## Abstract

α-Synuclein (αS), a protein that plays a central role in Parkinson’s disease and related synucleinopathies, is an intrinsically disordered protein (IDP) whose functional interactions and aggregation behavior can be strongly influenced by post-translational modifications (PTMs). Phosphorylation, acetylation, and other PTMs regulate αS’s interactions with lipid membranes and binding partners, whereas their dysregulation is associated with aggregation and neuronal toxicity. Despite significant progress through chemical and semi-synthetic approaches, investigating the combinatorial effects of PTMs has remained challenging due to the lack of accessible, site-specific methods. Here, we present an integrated strategy combining genetic code expansion, enzymatic modification, and intein-mediated click chemistry to generate αS variants bearing multiple defined PTMs and a C-terminal fluorescent label. The resulting constructs enable direct evaluation of how individual and combined PTMs influence αS structure, lipid binding, and cellular internalization. Our approach expands the molecular toolkit for dissecting PTM crosstalk in αS and other aggregation-prone IDPs, advancing mechanistic understanding and supporting the development of therapeutic strategies for neurodegenerative disease.

## Introduction

The protein α-Synuclein (αS) is a small protein (14.5 kDa) implicated in the pathogenesis of Parkinson’s disease (PD) and related diseases like multiple system atrophy (MSA), collectively termed synucleinopathies.^[1]^ Physiologically, αS is found primarily in the presynaptic terminals of neurons and its native roles are thought to be in vesicle trafficking and regulating neurotransmission.^[2] [3]^ αS has three domains: the positively charged N-terminal domain (residues 1-60), the hydrophobic Non-Amyloid Component (NAC) domain (residues 61-95), and the acidic C-terminal domain (residues 96-140).^[4]^ αS is intrinsically disordered in solution, and adopts an α-helical structure upon binding to lipid membranes.^[5]^ This conformational plasticity that is central to its physiological function also underlies its pathological tendency to form oligomers and fibrillar aggregates which are hallmarks of synucleinopathies.^[6] [7]^ Because this conformational landscape is highly tunable, post-translational modifications (PTMs) constitute a crucial regulatory mechanism for αS. PTMs regulate αS structure and function, modulating its membrane affinity, aggregation propensity, and protein – protein interactions.^[8]^ Among the two most common αS PTMs, constitutive N-terminal acetylation (AcNterm) prevents proteosomal degradation and promotes lipid association,^[9]^-^[9b]^ whereas phosphorylation at serine 129 (pS129) is a prominent modification in Lewy body aggregates in PD and thus is a hallmark of disease-associated αS.^[10] [11]^ Site-specific lysine acetylation (^Ac^K) has also emerged as a critical regulatory mechanism influencing αS’s charge distribution and conformational dynamics.^[12] [12b]^

Proteomic studies of human PD and MSA post-mortem patient samples have identified numerous αS PTMs, many of which co-occur.^[13]^ However, the combinatorial effects of these PTMs – potentially acting cooperatively or competitively – across the protein’s three domains remains poorly understood. Traditional semisynthetic and chemical ligation methods have enabled valuable site-specific insights but are technically demanding, low yielding, and inaccessible to most biochemistry or neurobiology laboratories.^[14] [15]^ This has created a major gap between conceptual advances in PTM mechanism and the experimental ability to study these modifications in a reproducible and combinatorial manner.

To bridge this gap, we have developed a robust, modular platform that utilizes genetic code expansion (GCE), enzymatic modification via co-expression, and intein-mediated click chemistry for the efficient generation of αS variants bearing multiple authentic PTMs as well as fluorescent labels. Our approach offers three major advantages: (i) it is fully compatible with standard *E. coli* expression workflows, (ii) it provides residue-level control of PTM combinations, and (iii) it produces sufficient quantities for both *in vitro* biophysical and biochemical studies, as well as cell-based assays. Beyond αS, this method provides a versatile platform applicable to other intrinsically disordered and aggregation-prone proteins, such as the Alzheimer’s disease associated protein tau, enabling systematic interrogation of PTM crosstalk mechanisms.

## Results and Discussion

The goal of this study was to establish a highly reproducible expression and labeling system capable of introducing multiple site-specific authentic PTMs into αS through recombinant protein expression (Figure 1). The platform utilizes four orthogonal plasmids encoding the following: (1) αS with a TAG codon and a C-terminal intein for functionalization and a 6x-His tag for purification; (2) the fission yeast N-terminal acetyltransferase NatB for N-terminal acetylation of αS; (3) pTECH-chAcK3RS, which encodes the aminoacyl tRNA synthetase (RS) and tRNA for GCE-based incorporation of *N*ε-acetyl-L-lysine (^Ac^K) at TAG sites in αS; and (4) PLK2 for selective phosphorylation of αS at serine 129. Cleavage of the intein removes the His-tag and, in the presence of select nucleophiles, allows for functionalization of the C-terminus; for example, sodium 2-mercaptoethane sulfonate (MeSNa) in combination with propargylamine generates an alkyne handle for click chemistry labeling, as described below.

**Figure 1.**
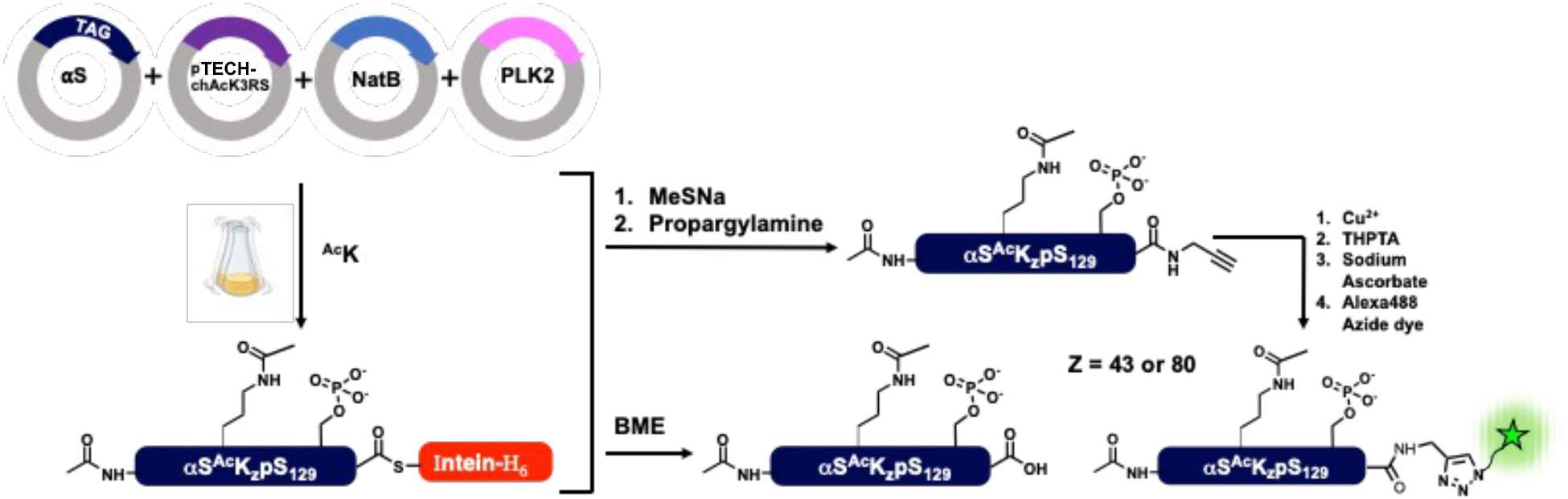
Approach used to generate modified αS. GCE, enzymatic co-expression and functionalization using selective intein cleavage followed by click chemistry to generate αS with three site-specific PTMs and a C-terminal fluorescent label.

The αS plasmid has been used previously with plasmids encoding NatB, chAcK3RS, and PLK2.^[7, 16]^ However, to use four plasmids in combination, they must have mutually orthogonal antibiotic resistance so that cells carrying all four plasmids can be selected, as well as compatible origins of replication to ensure propagation. Analysis of the four vectors used previously showed that pTXB1-αS, pTech-chAcK3RS, and pRSFduet-PLK2 were compatible, but the pACYCduet-NatB vector shared its antibiotic resistance (chloramphenicol) and origin of replication (p15A) with pTech-chAcK3RS (Table S1). We therefore cloned the NatB sequence into a pCDFduet vector with spectinomycin resistance and a CloDF13 origin of replication. Electrocompetent *E. coli* BL21(DE3) cells were co-transformed with the four plasmids encoding αS (with a TAG codon replacing either Lys43 or Lys80), the NatB acetyltransferase complex, the chAcK3RS/tRNA pair, and PLK2. Prior to inducing αS expression at OD_600_=0.6 with the addition of 1 mM IPTG, the media was supplemented with 10 mM ^Ac^K and 50 mM nicotinamide, an inhibitor of endogenous deacetylases. Following crude purification on a Ni-NTA column, αS was cleaved from the intein either in the presence 2-mercaptoethanol (BME) to generate a C-terminal carboxylic acid, or MeSNa followed by propargylamine treatment to introduce an alkyne handle. Following a second Ni-NTA column to remove cleaved 6x-His tagged intein, modified αS was purified by reverse phase high performance liquid chromatography (RP-HPLC) and its mass determined by matrix-assisted laser desorption ionization mass spectrometry (MALDI-MS), (Figure 2). Typical yields were 0.6 – 1 mg of purified αS per liter of culture with 100% site-selective incorporation of all intended PTMs.

**Figure 2.**
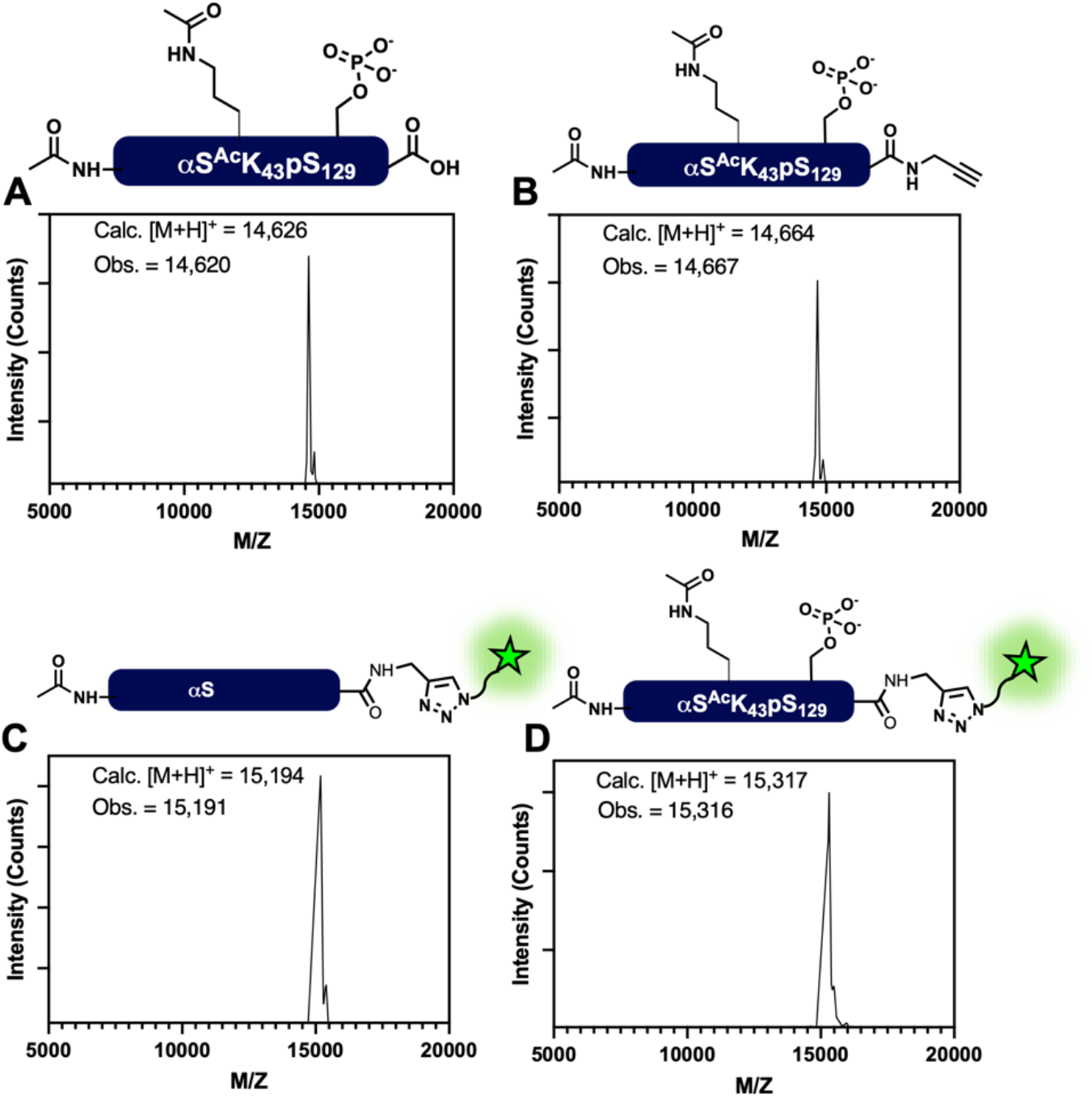
Characterization of αS-^Ac^Nterm^Ac^K_43_pS_129_ and labeled αS. **A**: MALDI-MS of protein following cleavage of the intein with BME, giving a C-terminal carboxylic functional group; **B** MALDI-MS of protein following MeSNa and propargylamine cleavage of intein to give a C-terminal alkyne handle. **C;** MALDI-MS of αS-^Ac^Nterm after labeling with Af488-N_3_. **D**; MALDI MS of αS-^Ac^Nterm^Ac^K_43_pS_129_ following Af488-N_3_ labelling.

The C-terminal alkyne handle allows for selective functionalization at a site that is not expected to perturb membrane binding or aggregation, given that the C-terminus of αS does not associate with membranes and remains disordered and unresolved in cryo-electron microscopy or solid state NMR structures of αS fibrils.^[17]^ Alexa Fluor 488 azide (Af488-N_3_) was conjugated to αS via copper-catalyzed click chemistry (See Supporting Information).^[18]^ The mass of the labeled protein, αS-Af488, was verified by MALDI-MS (Figure 2). Fluorescently labeled protein is required for a variety of different types of biophysical and biochemical assays, including single-molecule Förster resonance energy transfer (FRET) to measure conformational distributions^[5]^, fluorescence correlation spectroscopy (FCS) to quantify membrane binding^[19]^, cellular uptake and trafficking studies^[20]^ and aggregation kinetics^[21]^. Below we illustrate usefulness in lipid-vesicle binding assays, frequently used as a surrogate for αS-function, and spontaneous cellular uptake, of relevance to pathological spread.

### Vesicle Binding of αS-^Ac^Nterm^Ac^K_43_pS_129_

To determine the effect of modifications on interactions between αS and lipid membranes, we used FCS to quantify binding of αS-^Ac^Nterm^Ac^K_43_pS_129_ on to lipid vesicles.^[22]^ Lipid vesicles composed of 60:25:15 1-palmitoyl-2-oleoyl-glycero-3-phosphocholine/1-palmitoyl-2-oleoyl-*sn*-glycero-3-phosphoethanolamine/1-palmitoyl-2-oleoyl-sn-glycero-3-phospho-L-serine (POPC/POPE/POPS) were prepared by extrusion. Autocorrelation curves of αS-Af488 in buffer or in the presence of a high concentration of vesicles were initially measured independently to determine expected diffusion times for free protein and vesicles, respectively. To evaluate binding of αS to vesicles, a fixed amount of αS-Af488 was added to varying concentrations of vesicles and autocorrelation curves were measured. The fraction of vesicle-bound αS was determined by fitting the autocorrelation curves to a model for two fluorescent species, one corresponding to free αS-Af488 and the other to vesicle-bound αS-Af488. The fraction bound at each vesicle concentration plotted against the lipid concentration and fit with a hyperbolic binding model to determine an apparent dissociation constant *K*_d,app_. We compared αS-^Ac^Nterm and αS-^Ac^Nterm^Ac^K_43_pS_129_ and observed a more than 2-fold decrease in affinity for αS-^Ac^Nterm^Ac^K_43_pS_129_ (Figure 3). Previous studies in our laboratory have shown αS-^Ac^K_43_ to have weaker vesicle binding (∼20%) compared to wild type.^[23]^ In contrast, αS-pS_129_ has been reported to exhibit stronger binding to synaptic vesicles.^[24]^ Our results showed αS-^Ac^Nterm^Ac^K_43_pS_129_ has a decrease in affinity compared to αS-^Ac^Nterm, implying that acetylation at K_43_ overcomes the effect of phosphorylation at Ser_129_. This shows the importance of studying effects of PTMs combinatorially and hence the usefulness of this method.

**Figure 3.**
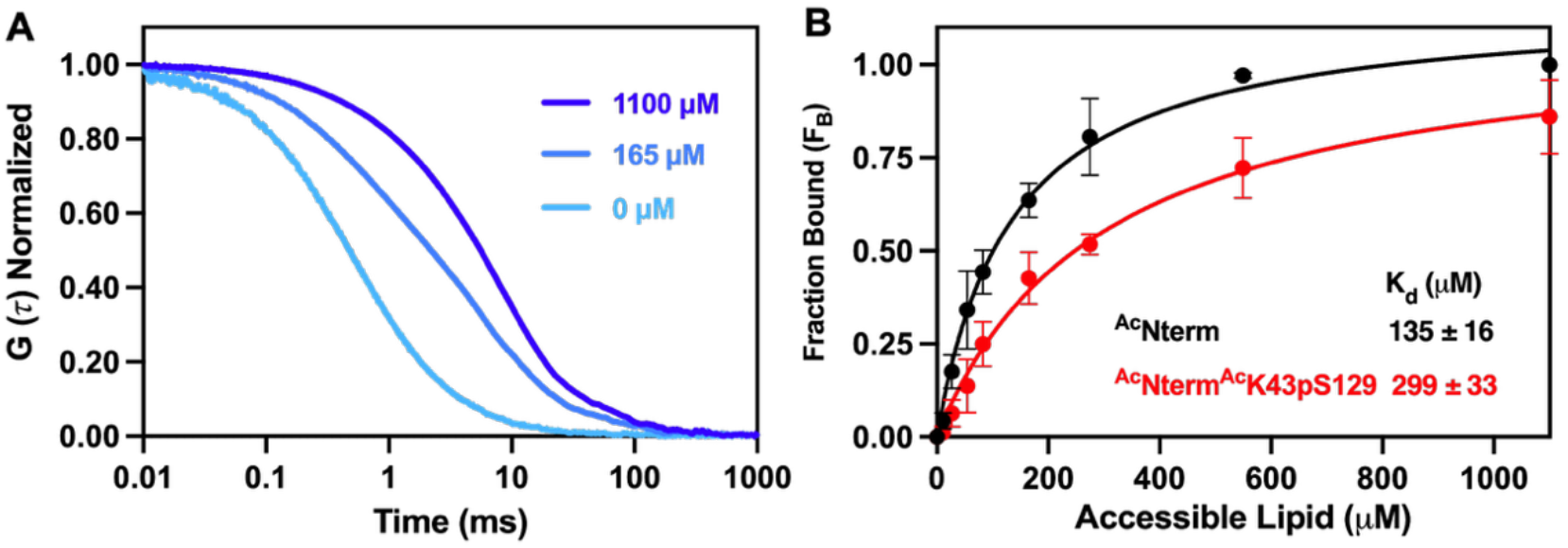
Quantification of αS binding to vesicles. (**A)** Representative autocorrelation curves for αS-Af488 with increasing vesicle concentrations (0, 165, and 1100 µM lipid). At higher vesicle concentrations, the curves decay at longer lag times, reflecting an increasing fraction of vesicle-bound αS. (**B**) Binding curves for αS-^Ac^Nterm and αS-^Ac^Nterm^Ac^K_43_pS_129_ demonstrate that the presence of the two PTMs decreases the binding affinity relative to αS-^Ac^Nterm.

### Cellular Internalization of αS-^Ac^Nterm^Ac^K_43_pS_129_

As mentioned above, uptake of extracellular αS may be relevant to pathological spread in disease and there is significant interest in characterizing the molecular details of the relevant pathways. As compared to more traditional immunofluorescence, which involves both fixing and labeling the cells with antibodies, fluorescently labeled proteins enable real-time visualization of protein localization and trafficking.^[25]^ To demonstrate that our strategy is compatible with such applications, αS-^Ac^Nterm^Ac^K_43_pS_129_ labeled with Af488 was incubated with SH-SY5Y neuroblastoma cells. Confocal imaging shows that the labeled protein was readily internalized and visualized in puncta indicative of endosomal processing (Figure 4).^[20]^

**Figure 4.**
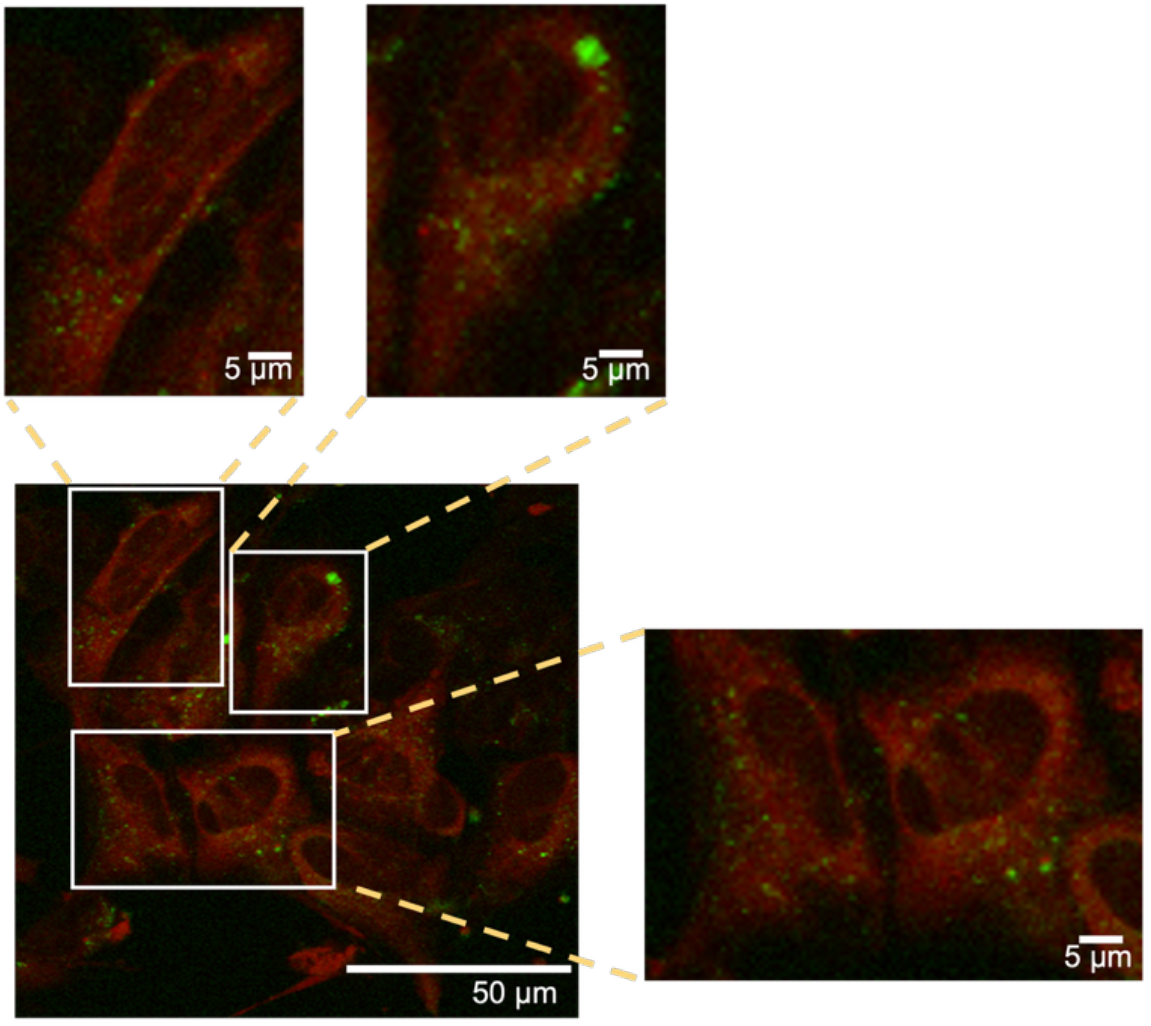
αS-^Ac^Nterm^Ac^K_43_pS_129_ labeled with Af488 is spontaneously internalized by SH-SY5Y cells. Representative time-resolved fluorescence microscopy images of SH-SY5Y cells after incubation with 200 nM αS-^Ac^Nterm^Ac^K_43_pS_129_ -Af488 (green, 482 nm excitation) for 16 h. Cell membranes were labeled with CellTracker Red CMPTX (red, 561 nm excitation). Images acquired at 37 °C, 5% CO_2_.

## Conclusion

In this work, we have described the production and characterization of αS with three authentic PTMs introduced by a combination of GCE and enzymatic modification. Additionally, the triply modified constructs were labeled at the C-terminus with Af488-N_3_ by copper-catalyzed click chemistry. These methods allow for site-specific incorporation of acetylation and phosphorylation at key sites in αS, while being broadly accessible to biochemistry laboratories that may lack peptide synthesis and ligation chemistry expertise. Additionally, we demonstrated the utility of these approaches for fluorescent labeling, allowing for biophysical or cell biology studies.

While this work focuses on αS as a model system, we envision that it could be applied to other disordered, aggregation-prone proteins in a straightforward manner. For example, co-expression of tau with the kinase GS3Kb yields tau phosphorylated at multiple sites^[26]^; inclusion of GCE could be used to introduce another site-specific PTM or a handle for bioorthogonal labeling with a fluorophore. In combination with the intein-cleavage mediated installation of the C-terminal alkyne group, this would allow for two orthogonal fluorescent labels for FRET measurements^[27]^ or spin-labels for NMR or electron paramagnetic resonance spectroscopy, while leaving tau’s two native cysteine intact and unmodified. Overall, this work showcases the value of GCE and enzymatic methods for producing PTM-modified proteins and their application in biophysical and structural characterization. Going forward, we will use this approach to investigate crosstalk between, and the combinatorial effects of, PTMs on function and dysfunction of this biomedically important class of proteins.

## Supporting Information

Descriptions of protein expression and modification, characterization of crude and purified proteins and experimental details for fluorescence spectroscopy and microscopy. The authors have cited additional references within the Supporting Information.^[5, 23, 28]^

## Acknowledgements

This research was supported by the University of Pennsylvania and the National Institutes of Health (NIH RF1NS125770 to E.R. and E.J.P.). Instruments supported by the NIH include MALDI-MS (NIH S10OD030460). B.A. was supported by a Hochstrasser Fellowship, M.S. was supported by a scholarship from the Nakajima Foundation and P. M-C. was supported by NIH T32 GM132039.

