## Supporting Information for "Genetic Code Expansion, Enzymatic Modification, and C-Terminal Labeling Enable Facile Production of Highly Modified α-Synuclein"

[b] Graduate Group in Biochemistry, Biophysics, and Chemical Biology
Perelman School of Medicine, University of Pennsylvania
206 Anatomy-Chemistry Building, 3620 Hamilton Walk, Philadelphia, PA 19104, USA

[c] Department of Biochemistry and Biophysics
Perelman School of Medicine, University of Pennsylvania
421 Curie Boulevard, Philadelphia, PA 19104, USA

### General Information

*E. coli* BL21(DE3) cells and *E. coli* Dh5α cells were purchased from New England Biotechnologies (Ipswich, MA, USA). DNA oligomers were purchased from Integrated DNA Technologies, Inc (Coralville, IA, USA). DNA extraction and Miniprep kits were purchased from Qiagen (Hilden, Germany). Buffers were made with Milli-Q filtered (18 MΩ) water (Millipore; Billerica, MA, USA). Preparation of the pTXB1-αS-intein$H_6$ plasmid containing α-synuclein (αS) with a C-terminal fusion to the *Mycobacterium xenopi* GyrA intein and C-terminal $His_6$ tag was described previously.[1] This plasmid was used as a starting point for the preparation of αS (mutants) -intein constructs with the primers given in Table S1. pTECHchAcK3RS (IPYE) was a gift from David Liu via Addgene (plasmid # 104069; http://n2t.net/addgene:104069; RRID:Addgene_104069; Watertown, MA, USA). Acetyllysine was purchased from ChemImpex (Wood Dale, IL, USA). Nicotinamide was purchased from Alfa Aesar (Tewksbury, MA, USA). AlexaFluor 488 azide was purchased from ThermoFisher Scientific. The lipids 1-palmitoyl-2 oleoyl-glycero-3-phosphocholine (POPC), 1-palmitoyl-2-oleoyl-*sn*-glycero-3-phosphoethanolamine (POPE) and 1-palmitoyl-2-oleoyl-*sn*-glycero-3-phospho-L-serine (POPS), were purchased from Avanti Research (Alabaster, AL, USA). Matrix-assisted laser desorption/ionization mass spectrometer (MALDI-MS) data were collected with a Bruker rapifleX MALDI-MS instrument or a Bruker Microflex MALDI-MS (Billerica, MA, USA). Ultraviolet-visible (UV-vis) absorption spectra were collected on a GENESYS 150 UV-vis spectrophotometer (Thermo Fisher Scientific; Waltham, MA, USA). Gel images were obtained with a Syngene G:Box mini-6 (Cambridge, UK). Proteins were purified on a 1260 Infinity II preparative high-performance liquid chromatography (HPLC) system (Agilent Technologies). Water + 0.1%

trifluoroacetic acid (TFA) (solvent A) and acetonitrile + 0.1% TFA (solvent B) were used as the mobile phase in HPLC.

**Generation of NatB plasmid**

The NatB subunits for incorporating N-terminal acetylation were cloned from the original pACYCduet vector into a pCDFduet vector to achieve orthogonal antibiotic resistance and origin of replication for all four plasmids used (Table S1).

| Target | Plasmid Name | Machinery Type | Antibiotic Resistance | Origin of Replication | Plasmid Compatibility Group* |
|---|---|---|---|---|---|
| **N-terminal acetylation** | pCDFduet | Enzymatic | Spectinomycin | CloDF13 | D |
| **Lys acetylation** | pTECH-chAcK3RS | GCE | Chloramphenicol | p15A | B |
| **Phosphorylation at S129** | pET28-PLK2 | Enzymatic | Kanamycin | ColE1 | A |
| **Alpha-Synuclein-GyrA** | pTXB1 | - | Ampicillin | ColE1 | A |

**Table S1**. Plasmid Compatibility Table
* Based on https://blog.addgene.org/plasmid-101-origin-of-replication and references therein.

The empty plasmid with backbone pCDFduet (Sigma, 71340-M) was a gift from Rick Cooley at Oregon State University. The pACYCduet-naa20-naa25 plasmid was a gift from Daniel Mulvihill at University of Kent, United Kingdom.

The following primers were designed:

**Table S2.** DNA Sequences for cloning

| | | |
|---|---|---|
| pCDF Backbone | Forward | 5′- GAACGCCAGCACATGGACTCGTCTACTAG -3’ |
| | Reverse | 5′- GATCCTGGCTGTGGTGATGATGGTGATG -3’ |
| NatB Backbone | Forward | 5’- CATCACCATCATCACCACAGCCAGGATC -3’ |
| | Reverse | 5’- CTAGTAGACGAGTCCATGTGCTGGCGTTC -3’ |

Deletion PCR was performed to amplify plasmid backbones and inserts with following conditions using standard PRC protocol. Upon completion of PCR, reactions were cleaned up with DNA Concentration & Purification kit from Zymo Research, then analyzed by agarose gel. For Gibson Assembly, NEBio calculator was used to calculate mass ratios of insert and vector. 100 ng vector, 2 equivalents of insert, nuclease-free water up to 10 µL, 10 µL 2x HiFi Assembly Master Mix were mixed and incubated at 50°C for 1h. The reaction was transformed into DH5-alpha cells, amplified in LB media with corresponding antibiotics and the DNA product was cleaned up with QIAprep Miniprep Kit.

**αS Expression Plasmids**

The following primers (Table S3) were designed for site-directed mutagenesis to introduce TAG (= Z) codons at each Lys acetylation site. Introduction of these TAG mutations to the pTXB1-αS-intein-$H_6$ plasmid has been described previously.[2]

**Primer Sequences:**

**Table S3.** DNA Sequences for cloning:

| | | |
|---|---|---|
| αS-$Z_{43}$ | Forward | 5'-GTAGGCTCCTAGACCAAGG -3' |
| | Reverse | 5'-ATAGAGAACACCCTCTTTTGTCTTTC -3' |
| αS-$Z_{80}$ | Forward | 5'- GCAGTAGCCCAGTAGACAGTGGAGGGA - 3' |
| | Reverse | 5'- TGTCACACCCGTCACCACTGC -3' |

**Expression and purification of αS**

To generate triply-modified αS, we followed the method previously described with modifications.[2] Plasmids encoding the following proteins (Table S1 for antibiotic resistance and Ori) were co-transformed by electroporation into BL21 (DE3) competent cells: αS fused to a His-tagged GyrA intein from *Mycobacterium xenopi* (αS-intein) and a TAG mutation at the site of interest; pTECH-chAcK3RS (IPYE) for incorporation of acetyl-lysine at the TAG codon; Plk2 (gift from Daniel T.S. Pak of Georgetown University) a kinase for site specific phosphorylation of serine 129,[3] and NatB for N-terminal acetylation.[4] Cells grown on ampicillin/chloramphenicol/kanamycin/streptomycin (Amp/Chlor/Kan/Strep) plates. Single colonies were picked to inoculate primary cultures in LB media supplemented with 0.1 mg/mL Amp, 0.034 mg/mL Chlor, 0.025 mg/mL Kan and 0.1 mg/mL Strep. Primary cultures were incubated at 37 °C overnight or until they were cloudy. Secondary cultures in LB media with antibiotics were inoculated and grown at 37 °C with shaking at 250 rpm until the optical density reached ~ 0.6 and then cooled to 18 °C. 50 mM nicotinamide and 10 mM Nε-acetyl lysine were added to the culture and incubated for ~ 5 min before inducing the expression of the gene of interest with 1 mM isopropyl ß-D-1- thiogalactopyranoside (IPTG). To generate αS with only N-terminal acetylation, protein expression was performed as above with the following differences: (1) cells co-transformed only with the NatB plasmid and αS-intein lacking any TAG codons; (2) nicotinamide and Nε-acetyl lysine were not added; (3) cells were plated and incubated on ampicillin/chloramphenicol (Amp/Chlor) plates.

Following overnight growth at 18 °C, the culture was centrifuged at 4600x g for 20 minutes at 4 °C. Cell pellets were re-suspended in buffer (40 mM Tris pH 8.3, a Roche protease inhibitor tablet, and 1 mM phenylmethylsulfonyl fluoride). Cells were lysed by sonication in an ice bath, (5 min, 1 s ON, 2 s OFF) and the resulting lysate was centrifuged at 20,000x g for 30 min at 4 °C to remove cellular debris. The supernatant (after centrifugation) was filtered with a 0.22 µM syringe filter and added to 5–7 mL of Ni-NTA resin equilibrated with αS-intein buffer 1 (ASB1; 50 mM HEPES pH 7.5), followed by incubation with rocking at 4 °C for ~1 hour. The resin was then washed with ~15 mL ASB1, followed by a wash with αS-intein buffer 2 (ASB2; 50 mM HEPES pH 7.5, 5 mM imidazole). The protein was eluted with ~12 mL Ni-NTA αS-intein buffer 3 (ASB3: 50 mM HEPES pH 8, 400 mM imidazole). Cleavage of the intein was carried out by incubation with 200 mM βmercaptoethanol (βME) overnight at room temperature on a rotisserie to give a c-terminal carboxylic group. Cleaved αS was dialyzed into 20 mM Tris, pH 8 buffer before purification over a second Ni-NTA column to remove the free intein and any remaining uncleaved αS-intein from the sample. The αS protein was purified by RP-HPLC over a C4 column, dialyzed into 1x phosphate buffered saline (PBS) pH 7.4, concentrated in an Amicon centrifugal filter, aliquoted and flash frozen for storage at -80 °C.

**One-Pot Cleavage of αS-Intein with MeSNa and Propargylamine**

To generate αS constructs with a propargyl amide at the C-terminus, αS was purified as described above through the elution from the Ni-NTA resin. The intein was cleaved by incubation with 150 mM MeSNa on a rotisserie overnight at room temperature. Following the overnight incubation, 300 mM propargylamine was added and the solution was incubated for an additional 5 hours. The solution was dialyzed into 20 mM Tris pH 8 buffer before purification over a second Ni-NTA column to remove the free intein from the cleaved αS protein. The αS protein was purified by RP-HPLC over a C4 column dialyzed into 1x phosphate buffered saline (PBS) pH 7.4 and concentrated in an Amicon centrifugal filter, aliquoted and flash frozen for storge at -80 °C until further use.

**Fluorescent Labeling of αS**

αS protein stocks in 20 mM 1x PBS were fluorescently labeled via copper-catalyzed azide-alkyne cyclization. A catalytic mixture consisting of 2 equiv $CuSO_4$, 10 equiv tris(3-hydroxypropyltriazolylmethyl)amine (THPTA), and 20 equiv sodium ascorbate was incubated at room temperature for 10 min and added to the protein along with 2 equiv Alexafluor 488 azide fluorophore (Af488-$N_3$). The mixture was degassed, and the reaction tube was wrapped with aluminum foil and incubated at 50 °C in a water bath for 10 mins, followed by 20 min incubation at 30 °C with shaking until product formation was observed by MALDI-MS. The labeled protein was exchanged into 20 mM 1x PBS, pH 7.4 and unreacted dye was removed by passing the solution over two coupled HiTrap Desalting columns. Following purification, the sample concentration was determined using a NanoDrop™ spectrophotometer using the following extinction coefficients: Af488 ε@494 nm = 73000 $M^{-1}cm^{-1}$, αS ε@280 nm = 5960 $M^{-1}cm^{-1}$ and calculated by correcting the absorbance signal at 280 nm for absorbance by Af488 using the following equation: A(αS)280 = [A280-0.11(A494)]/A280]. The protein was then aliquoted into microcentrifuge tubes, flash frozen, and stored at -80 °C.

**Preparation of Synthetic Vesicles**

Large unilamellar vesicles were prepared by extrusion through porous membranes. A mixture in 60:25:15 molar ratio of 1-palmitoyl-2-oleoyl-glycero-3-phosphocholine: 1-palmitoyl-2-oleoyl-*sn*-glycero-3-phosphoethanolamine: 1-palmitoyl-2-oleoyl-*sn*-glycero-3-phospho-L-serine (POPC: POPE: POPS) were drawn from chloroform stock and dried under nitrogen gas to form a film inside a glass vial. Films were desiccated under vacuum and re-hydrated in vesicle buffer (10 mM 4-(2-hydroxyethyl)piperazine-1-ethanesulfonic acid (HEPES), 150 mM NaCl, pH 7.4). Ten freeze-thaw cycles consisting of cooling in liquid nitrogen for 40 s and warming in a 60 °C water bath for 2 min were performed to aid the formation of uniformly sized vesicles. The aqueous lipids solution was then extruded 31 times through two stacked 50 nm pore membranes (Cytiva, Marlborough, MA, USA) in a Liposofast extruder (Avestin, Ottawa, ON, Canada). Vesicles were determined by to be monodisperse and distributed uniformly around 90 nm in diameter by dynamic light scattering (DLS), consistent across different concentrations of all samples. DLS data were collected on the ZetaSizer Nano Series Nano-ZS (Malvern Panalytical, Westborough, MA). All lipid vesicles were prepared fresh and used within 48 h of extrusion.

**Fluorescence Correlation Spectroscopy (FCS)**

Fluorescence correlation spectroscopy (FCS) measurements used to quantify binding of αS to lipid vesicles, were carried out as described previously [2, 5][6] . Eight-well chambered cover glasses (Nunc, Rochester, NY, USA) were prepared by plasma cleaning followed by incubation overnight with polylysine-conjugated polyethylene glycol (PEG-PLL). PEG-PLL coated chambers were rinsed with and stored in Milli-Q water until use. FCS measurements were performed on a lab-built instrument based on an Olympus IX71 microscope with a continuous emission 488 nm DPSS 50 mW laser (Spectra-Physics; Santa Clara, CA, USA). The laser power entering the microscope was adjusted to 4.5 μW. Fluorescence emission collected through the objective was separated from the excitation signal through a Z488rdc long pass dichroic filter and an HQ600/200m bandpass filter (Chroma; Bellows Falls, VT, USA). Emission signal was focused onto the aperture of a 50 μm optical fiber coupled to an avalanche photodiode (Perkin Elmer; Waltham, MA, USA). A digital autocorrelator (Flex03Q-12, correlator.com; Bridgewater, NJ, USA) was used to collect 10 autocorrelation curves of 10 seconds for each measurement of free protein in buffer without lipids or 30 autocorrelation curves of 30 seconds for each measurement in the presence of lipid vesicles. Global fitting was done using lab-written coded in GraphPad Prism version 10 (GraphPad Software; San Diego, CA, USA).

To determine the diffusion time of the proteins, each αS variant labeled with Af488 was measured in buffer. The average of 10 autocorrelation curves was fit to a 1-component autocorrelation function in GraphPad Prism:

$$G(\tau) = \frac{1}{N}\left(\frac{1}{1+\frac{\tau}{\tau_1}} * \left(\frac{1}{1+\frac{s^2\tau}{\tau_1}}\right)^{\frac{1}{2}}\right) \qquad \text{(S1)}$$

where G($\boldsymbol{\tau}$) is the autocorrelation function, N is the number of molecules in the focal volume, $\boldsymbol{\tau}_1$ is the diffusion time of αS, and s is the radial-to-axial ratio of the excitation volume.

**Calculation of Vesicle Binding Affinity**

Autocorrelation curves of αS - labeled with Af488 were examined in the presence of varying concentrations (0.011 mM to 2.0 mM lipid) of lipid vesicles consisting of 60:25:15 POPC/POPE/POPS. The 30 curves were averaged together to obtain statistical variation, which used for weighting in the fitting to an equation with a two diffusing species:

$$G(\tau) = \frac{1}{N}\left(A * \frac{1}{1+\frac{\tau}{\tau_1}} * \left(\frac{1}{1+\frac{s^2\tau}{\tau_1}}\right)^{\frac{1}{2}} + Q * (1-A) * \frac{1}{1+\frac{\tau}{\tau_2}} * \left(\frac{1}{1+\frac{s^2\tau}{\tau_2}}\right)^{\frac{1}{2}}\right) \qquad \text{(S2)}$$

where G($\boldsymbol{\tau}$) is the autocorrelation function, N is the number of molecules in the focal volume, $\boldsymbol{\tau}_1$ is the diffusion time of αS, $\boldsymbol{\tau}_2$ is the diffusion time of the vesicles, s is the radial-to-axial ratio of the excitation volume, Q is the ratio of the brightness of vesicle-bound αS relative to αS, and A is the fraction of free αS. Diffusion times for $\boldsymbol{\tau}_1$ were constrained to the values obtained for each αS construct measured independently and fit to a single-species diffusion model as described above. Diffusion times for $\boldsymbol{\tau}_2$ were constrained to the maximum diffusion time observed on that experimental day, also determined from single-species model fits with $\tau_2$ set as a global parameter in GraphPad Prism in Equation S2. As with the Eqation S1, s was held constant. All remaining parameters (N, Q, and FF) were free parameters. The fraction of bound αS at each lipid concentration was used to generate a binding curve which was fit to the following equation:

$$D = \frac{B_{max}X}{K_{d,app}+X} \qquad (S3)$$

Where D is the fraction of αS bound, x is the accessible lipid concentration, $B_{max}$ is the maximum fraction of αS bound, and $K_{d,app}$ is the apparent dissociation constant. Averages and standard deviations were calculated from at least 3 independent measurements performed on separate days at each lipid concentration.

**Cell uptake of αS**

SH-SY5Y cells (ATCC, Cat. CRL-2266) were maintained in T25 flasks at 37 ºC in a humidified incubator with 5% CO2 in DMEM F12 media (Gibco, Cat. 11320033) supplemented with 10% Fetal Bovine Serum (FBS) and 1% Penicillin/Streptomycin (P/S). 8-well ibidi chambers (μ-Slide, 8-well, ibiTreat, ibidi GmbH, Germany) were coated with Poly-L-lysine and seeded with 80,000 cells/well. Upon reaching 80-90% confluency, cells were exchanged into uptake media [phenol free DMEM F12, 1% P/S] supplemented with 200 nM αS-Af488 and incubated for 16 hours. The uptake media was exchanged for membrane staining media [phenol free DMEM F12, 1% P/S, 1 μM CellTracker Red CMPTX dye] (Invitrogen, Cat. C34552) and incubated for 30 minutes and then were washed three times and left in imaging media [phenol free DMEM F12, 1% P/S, 10% FBS]. Images were taken on a PicoQuant MicroTime 200 Time-resolved Fluorescence Microscope equipped with a Flimbee galvo scanner (PicoQuant, Berlin, Germany). The cells imaged in a portable stage-top incubator (Tokai Hit, STRF-WELSX-Set) maintained at 37 ºC and supplied with 5% $CO_2$. For imaging the αS-Af488, a 482 nm laser with a 525 35 nm band pass filter was used; for the cell stain, a 561 nm laser with a 600 nm long-pass filter was used. Single frame images were acquired at a resolution of 512 x 512 pixels with a dwell time of 60 μs. Images were exported as TIFF files and analyzed using Fiji ImageJ software by adjusting the intensities of the images to improve visualization of αS-Af488 uptake.

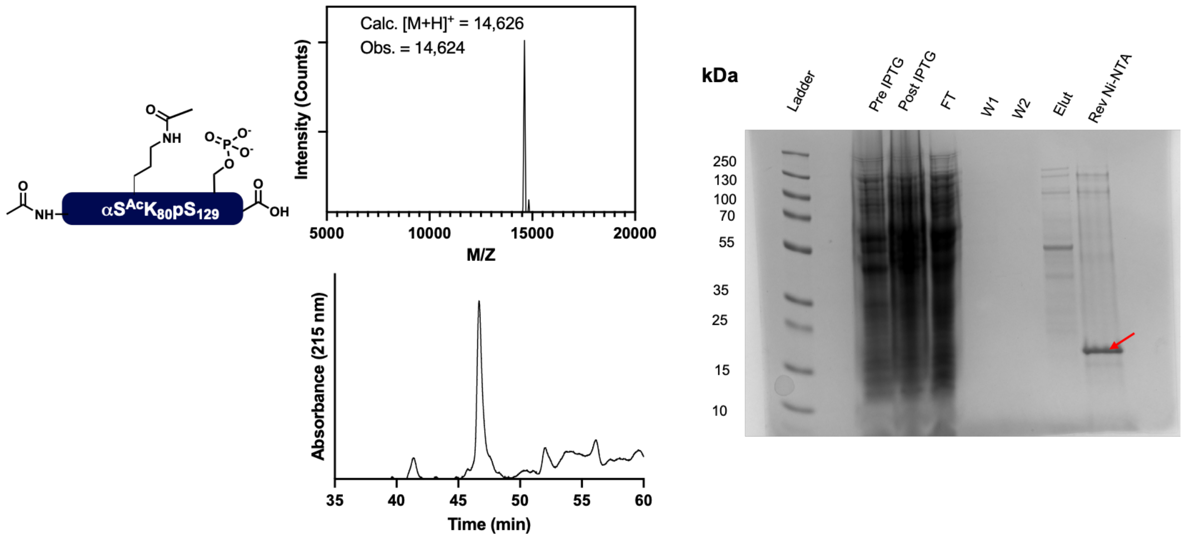


**Figure S1**. Gel, HPLC, and MALDI-MS analysis of αS- $^{Ac}$Nterm$^{Ac}$K$_{80}$pS$_{129}$ from genetic code expansion and enzymatic co-expression, followed by cleavage with BME.

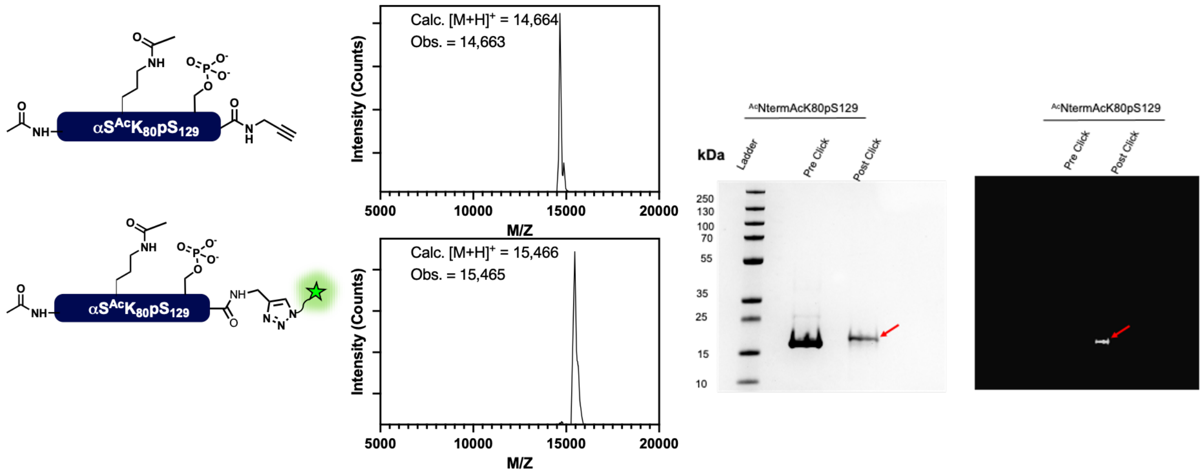


**Figure S2.** Gel, HPLC, and MALDI-MS analysis of αS-$^{Ac}$Nterm$^{Ac}$K$_{80}$pS$_{129}$$^{Atto488}$ from genetic code expansion and enzymatic co-expression and cleaved with MeSNa and propargylamine and labeled with an Atto488 azide dye via copper-catalyzed click chemistry.

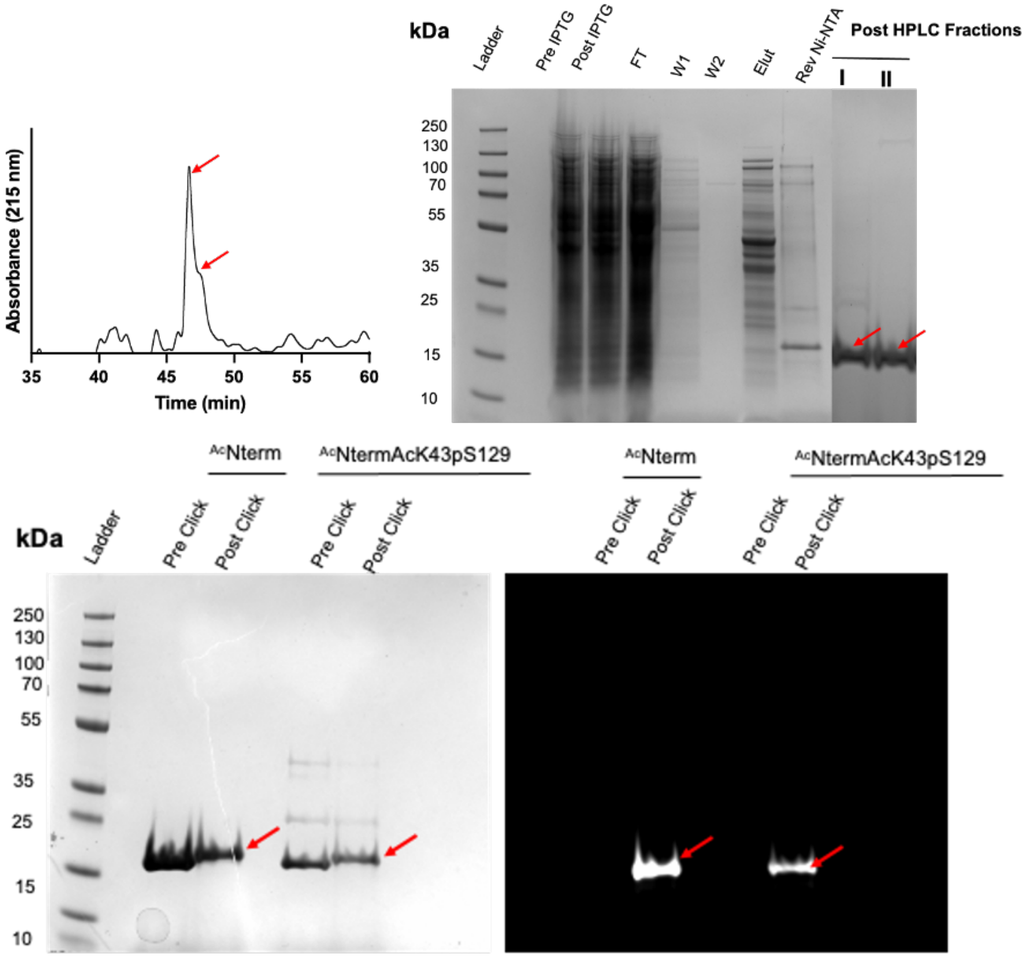


**Figure S3.** HPLC and Gel of $\alpha$S-$^{Ac}$Nterm$^{Ac}$K$_{43}$pS$_{129}$ cleaved with MeSNa and propargylamine (top) and Gels of $\alpha$S-$^{Ac}$Nterm$^{Af488}$ and $\alpha$S-$^{Ac}$Nterm$^{Ac}$K$_{43}$pS$_{129}$$^{Af488}$ (bottom).

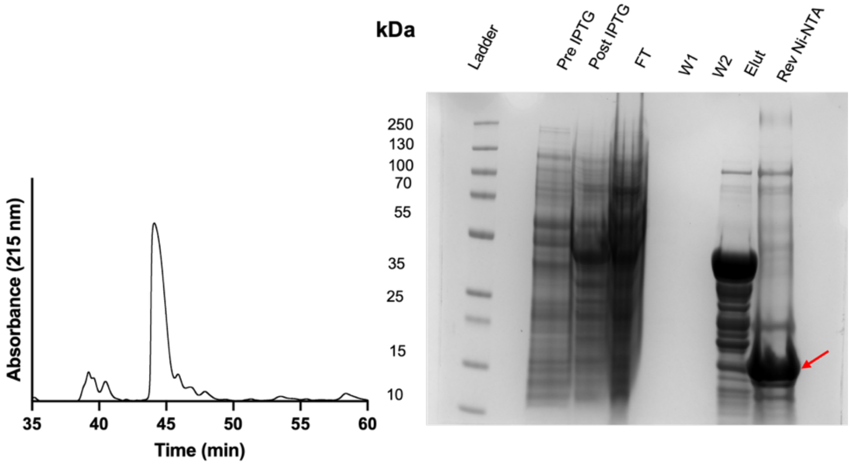


**Figure S4.** Gel and HPLC analysis of αS-$^{Ac}$Nterm construct.

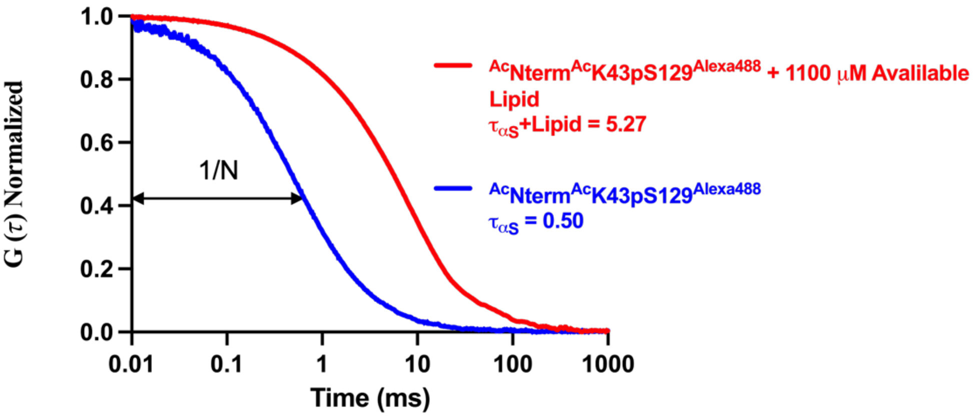


**Figure S5.** Representative autocorrelation curves of αS-$^{Ac}$Nterm $^{Ac}K_{43}pS_{129}^{Af488}$ (N= 9.7) and αS-$^{Ac}$Nterm $^{Ac}K_{43}pS_{129}^{Af488}$ +1100 μM Available Lipid (N = 36.7); curves were normalized to 1/N to allow for comparison.
